# Biodiversity conservation in cities: Defining habitat analogs for plant species of conservation interest

**DOI:** 10.1101/704700

**Authors:** M Itani, M. Al Zein, N. Nasralla, S. N. Talhouk

**Affiliations:** Department of Landscape Design and Ecosystem Management, Faculty of Agricultural and Food Sciences, American University of Beirut, Lebanon; Department of Biology, Faculty of Arts and Sciences, American University of Beirut, Lebanon; Nature Conservation Center, American University of Beirut, Lebanon

**Author notes:** Corresponding author (SNT).

## Abstract

Urban plant habitats have become primary drivers of species interactions. They consist of managed vegetation and spontaneous assemblages of native, naturalized, ornamental garden escapes, and invasive species. Our objective was to define urban habitat analogs for a plant species of conservation interest, *Matthiola crassifolia,* which has persisted in varying abundance in the Mediterranean city of Beirut.

We adopted a stepwise method that integrates two vegetation assessments, floristics, and physiognomy. We placed seventy-eight quadrats (1m x 1m) in 12 study sites following a deliberate biased method to capture habitat diversity. In every quadrat, we performed taxonomic identification and recorded life form of each species. We pooled species that shared the same life form into categories and estimated area cover for each of these life forms. We performed TWINSPAN analysis on floristic data to identify species positively associated with *M. crassifolia,* and on life forms, to determine plant assemblages that promote optimal *M. crassifolia* representation. We then combined findings from both analyses to generate a description of urban habitat analogs suitable for *M. crassifolia*.

The results revealed that urban habitat analogs favorable to *M. crassifolia* include green spaces dominated by palms, low-lying succulents, or by shrubs with scale-like leaves. On the other hand, spaces dominated by turf grass, canopy trees, or vegetation that produces significant litter were not favorable to *M. crassifolia*’s persistence. Based on these findings, we generated a plant palette of native and non-native species to design urban habitat analogs favorable to the persistence of *M. crassifolia*.

**Synthesis and applications:** The application of this method can inform planting designs that yield suitable habitats for plants of conservation interest. It can also guide landscape management plans that seek to create or modify green spaces to optimize growing conditions for species of conservation interest. Depending on sites, and based on the information generated by the stepwise method, designers and managers may decide to exclude life forms of native or non-native species that do not support the growth of a species of conservation interest, or they may create an artificial habitat that is conducive to its persistence.

## Introduction

Ornamental, native, naturalized, garden escapes, and invasive plant species, grow in managed, partially managed or unmanaged artificial urban habitats. Native plant species of conservation interest can adapt to such disturbed urban conditions depending on their ruderal behaviors [1]. However, their persistence is unlikely as their fate depends on how these artificial habitats are conceived, designed, and managed. This becomes critical when the geographic distribution of a species lies within the boundaries of the city. Urban biodiversity strategies have proposed to transform artificial urban habitats into habitats suitable for native plant conservation [2]. One example of urban biodiversity strategy is the use of species-rich herbaceous communities to promote biodiversity in cities [3]. Another strategy, referred to as reconciliation ecology, proposes the conversion of spaces assigned to human activities into spaces that support the persistence of native species [4]. Identifying habitat analogs in this case is essential to guide reconciliation ecology strategy in cities [5]. Provided appropriate conservation targets, habitat analogs could dilute the distinction between disturbed and non-disturbed habitats as favorable sites for plant conservation [6, 7]. Collecting data to inform and guide urban biodiversity strategies is challenging because all currently available methods are intended for field studies in natural areas and do not always yield clear findings in urban contexts (Table 1).

**Table 1.** Methods used to describe vegetation.

| Technique /<br>reference | Type | Description of method | Presentation of<br>results | Reference |
| --- | --- | --- | --- | --- |
| Raunkiaer's life<br>form classification | Physiognomic | All species in a study area categorized into Raunkiaer life forms of which five main categories were described by Raunkiaer himself on basis of the height of perennating buds above the ground. | Bar graph for<br>identified life<br>forms showing<br>percent species | [8, 9, 10, 11] |
| Dansereau's method<br>(1951, 1957) | Physiognomic and<br>structural | Structure of vegetation described based on six criteria sets including life form, size, cover, function and leaf size, shape and texture | Symbolic profile<br>diagrams of site | [12, 13] |
| Kuchler's method<br>(1967) | Physiognomic and<br>structural | Subdivides site vegetation into a hierarchy starting with woody and herbaceous categories further divided into seven and three classes respectively. Each physiognomic class is further described on bases that include leaf characteristics, height and cover. | Formulae that<br>employ letters and<br>numbers | [14] |
| Fosberg's method<br>(1961) | Physiognomic and<br>structural | Subdivides site vegetation into a four level hierarchy. The vegetation is first categorized as open, closed or sparse. At the second level, 31 formation classes are recognized based on height and continuity. The third and fourth levels describe plant function and leaf and growth form of dominant species respectively. Mapping of formation groups is conducted at the last level. | Formulae that<br>employ letters and<br>numbers | [15] |
| Habitat classification | Structural | Divides habitat systems into categories. Terrestrial habitat system is subdivided into four categories. Based on height of dominant vegetation, habitat of a site is categorized as open ground, field layer or woodland. Ecosystem structure and diversity is assessed after measurements of area covered by each habitat type are obtained. Number of species is counted by using sample quadrats in each habitat type for comparison purposes. | Lists of groups of sites representing main habitat types in an area that can be mapped | [16, 17] |
| Community classification | Floristic | Plants occurring in each sampling quadrat are identified to the species level and the abundance of each is measured. Floristic data is used to classify quadrats into groups based on how similar they are. | Lists of species and quadrat groups representing communities | [18] |
| EcoVeg | Physiognomic, floristic and Ecological | Follows hierarchal classification based on a set of vegetation criteria in conjunction with ecological characteristics and according to whether vegetation is natural or cultural, the method follows different rationales. | Upper, mid and lower levels subdivided into various classification | [19] |

For example, studies that used floristics to describe urban habitats found an over-representation of ruderal species and high taxonomic diversity between relatively close sites [18, 20, 21, 22, 23, 24, 25, 26, 27, 28, 29, 30, 31, 32, 33, 34, 35]. In contrast, physiognomic and structural vegetation description, developed to describe natural vegetation over large areas, may be a more useful tool for urban biodiversity than floristics. Physiognomy reflects predominance of life strategies adopted by different life forms and it is applicable in highly modified sites and at both macro- and micro-climate conditions [36, 18, 37].

In addition to field assessment challenges in cities, the success of plant conservation strategies is also highly influenced by social perception and preference and should take into consideration such requirements. For example, studies have shown that spontaneous ‘unmanaged’ vegetation may not appeal to residents as aesthetically pleasing nor is it perceived as acceptable ‘urban nature’ by decision-makers [38, 39, 40]. This is further complicated by the fact that plant selection and management, is driven by landscape architects and landscape contractors who have limited experience with native species, and do not have clear guidelines to contribute to biodiversity conservation in cities [41, 42].

The objective of this study was to define urban habitat analogs for a plant species of conservation interest, *Matthiola crassifolia,* which has persisted in varying abundance in the Mediterranean city of Beirut.

## Materials and methods

### Species of conservation interest and its distribution

Named after Pietro Andrea Mattioli, *Matthiola* R.Br. is a widespread genus of flowering plants represented by about 48 species ranging from annual, biennial and perennial, woody and herbaceous plants and sub-shrubs, many of which are heavily scented, and available in a wide range of colors for horticulture and floristry [43]. The genus *Matthiola* can be split into 12 distinct species-groups [44] with species distributed throughout Macaronesia, the Mediterranean basin, the Saharo-Sindian region and NE Africa–Asia, and it exhibiting two centers of taxonomic diversity in Turkey and the Irano-Turanian region.

There are four *Matthiola* species recorded in Lebanon, two of which are either national or regional endemics. The Species-Group OVATIFOLIA is represented by the regional endemic *Matthiola damascena* Boiss. The Species-Group LONGIPETALA is represented by *Matthiola tricuspidata* and *Matthiola longipetala.* Species-Group INCANA is represented by the national endemic *Matthiola crassifolia* Boiss. & Gaill which is restricted to a few locations along the highly urbanized Lebanese coast and is the subject of this study. *M. crassifolia* is a taxon of conservation interest as the species is recognized as an endemic of Lebanon. However, [44] has questioned the taxonomic status of the species proposing that it be considered subspecies of *Matthiola sinuata*. Even if future molecular analyses support this preference, the taxon will remain an endemic of Lebanon yet at the intra-specific level.

The most comprehensive record of the distribution and status of *M. crassifolia* prior to this study was by [45] who performed a systematic survey of the Lebanese coast and recorded the presence of the species in three out of five previously reported sites, Beirut and Byblos. Subsequent field investigations by [46] added Saida, Khaldeh and Amchit as localities for *M. crassifolia*. Our field survey to these localities confirmed the extinction of *M. crassifolia* in Saida and its continued presence in Khaldeh, Beirut and Byblos [47].

### Study area

Beirut (33.8869° N, 35.5131° E), the capital of the Republic of Lebanon, is located on the eastern coast of the Mediterranean. Archeological evidence shows that humans have continuously occupied Beirut for the last 5000 years [48, 49]. Today, Beirut has one of the highest urban densities in the Middle East with an area roughly over 20 km^2^, population density is estimated at 21,000 people per sq. km [50 51]. The topography of the city includes two hills, Achrafieh (100 m elevation from the sea) and Mousseitbeh (80 m elevation from the sea) [52]. Paul Mouterde, who conducted floristic studies in Beirut in the 20th century, reported 1200 floral species including native, naturalized, ornamental garden escapes, and invasive species [53].

Ras Beirut, our study site, is defined by a 6 km long and 2 km wide cape [54]. Today, this area consists of densely populated neighborhoods interspersed with managed landscapes and zones with spontaneous naturalized vegetation occurring within geographically adjacent lots. Recent floristic studies of semi natural areas in Ras Beirut revealed low community similarity, patchy species distribution, and predominance of habitat non-specific species [55]. Green spaces in Ras Beirut fall under two broad categories; managed landscapes, dominated by exotic ornamental species planted in raised beds with reconstructed soil, and spontaneous landscapes where spontaneous floral communities survive along with naturalized garden escapees, in coastal cliffs, along the rocky water front, and in un-built/abandoned lots [56]. Following early botanical studies of semi natural areas in Beirut, the city has been subjected to extensive landscape transformation, and today it still harbors a significant remnant native vegetation. Based on these facts, Beirut can be considered as a type-three city that is likely to be carrying an extinction debt [57].

### Field data collection

We used a deliberate biased method to select study locations and to lay out sampling quadrats [58]. We set a total of 78 quadrats in 12 sites. We placed quadrats, 1 m × 1 m, in anthropogenic habitats and in semi natural habitats that do not include shrubby vegetation [18]. We placed larger quadrats, 2 m × 2 m, in locations where shrubs are present [18]. As in Dinsdale, deliberate bias method consists of placing quadrats in areas judged representative of the selected location and for capturing the maximum observed variation [58, 59].

We made three modifications to the sampling technique to address site-specific issues; 1) When the boundary of a given plant community was not clearly defined due to site disturbance, we set quadrats within assumed boundaries of the community to capture plant diversity, 2) In cases where species had an ‘individualistic’ distribution pattern adding to the difficulty in conceiving boundaries [60], we increased the number of quadrats to capture the observed variation, 3) Since we do not know the dispersal distance of the target species, when a vegetation community harbored the target species we placed two quadrats; we set one to include the target species and placed the other quadrat in a location where the target species did not grow. In communities that did not harbor the target species, we set only one quadrat.

We divided each quadrat into a grid of 100 subunits to ensure speed of measurement and relative accuracy [61, 62, 63, 64]. In every quadrat, we determined percent cover using the 11-point Domin cover scale by visually assessing subunits as: fully covered, empty, and partially covered for each species and each life form [18]. Data obtained from all subunits within a quadrat was then added to determine Domin cover per quadrat.

### Taxonomic and life form identification

We identified each plant specimen by consulting published floras, voucher specimens at the University Herbarium (Post Herbarium), and photographic floras [65, 53, 46]. All identified species were described by their life form according to Ellenberg and Mueller-Dombois amended to include bunched shoot arrangement in reptant hemicryptophytes which forms a partially decomposed thick mat and peat accumulation [10]. We then pooled species that shared the same life form under the one category and estimated area cover for each life form accordingly.

### Analysis

Based on the 11-point Domin cover scale, we analyzed floristic data, species and percent species cover, using TWINSPAN [66]. Using the same tool, TWINSPAN, we analyzed the life form data, which included life-form categories and percent cover as relative abundance of each life form within each quadrat. In the TWINSPAN, the cut levels 0-3-4-5-6-8 were applied. The TWINSPAN groups were characterized by constancy-percentage, average cover and representation of target species. A matrix was created to find intersections between quadrat groups defined by classifying life form and floristic data sets. This process led to the identification of new quadrat groups that share similar life form and species composition. The full dataset can be found in [47].

## Results

*M. crassifolia* is most widely distributed in Beirut; based on our field surveys its presence was confirmed in 73 sites of which only one site, Pigeon Rock (Site 17), is protected by law, and another site, the limestone cliff facing Pigeon Rock (Site 16), is almost inaccessible and may be considered *de facto* protected. The remaining 71 sites offer highly diverse habitats and are not protected [47]. In remnant semi-natural sites *M. crassifolia* is found in spiny Mediterranean heaths, screes, sea cliffs and rocky offshore islands, growing on both sandstone and limestone formations and on (stabilized) coastal sand dunes. In anthropogenic sites, it grows near open sewers, in abandoned dump sites, through cracks in concrete walls and asphalt, on heaps of gravel, in street medians and on two occasions, almost epiphytically, out of the trunks of date and fan palms. The species’ tendency to utilize modified habitats reflects its partial behavior as a ruderal [36]. Over the course of the study, *M. crassifolia* was lost in 20 sites to urban development including one site which harbored the largest clump counts. Only four of these sites were recolonized during the course of the study. The extent of this species in Beirut was reduced by 800 m as a conserquence of this loss which accounts for a decrease of 17% in the plant’s range in the city over a period four years [47].

We recorded the presence of 124 plant species belonging to 40 families and 107 genera in the 78 sampled quadrats [47]. Plant species co-occurring with *M. crassifolia* shown in Table 2 include 16% non-native species (Table 2).

**Table 2.** Plant species co-occurring with Matthiola crassifolia in Beirut (* non-native species).

|  |  |  |
| --- | --- | --- |
| *Aegilops geniculata* Roth | *Erigeron bonariensis* L.* | *Phleum subulatum* (Savi) Asch. & Graebn. |
| *Agave americana* L.* | *Erigeron canadensis* L.* | *Phyla nodiflora* (L.) Greene* |
| *Agave attenuata* Salm-Dyck* | *Euphorbia terracina* L. | *Picris rhagadioloides* (L.) Desf. |
| *Alcea setosa* (Boiss.) Alef. | *Ficus carica* L. | *Piptatherum miliaceum* (L.) Coss. |
| *Alyssum strigosum* Banks & Sol. | *Ficus microcarpa* L.f.* | *Pittosporum tobira* (Thunb.) W. T. Aiton* |
| *Amaranthus hybridus* L.* | *Galium canum* DC. | *Plantago coronopus* L. |
| *Ambrosia maritima* L. | *Galium murale* (L.) All. | *Plantago lagopus* L. |
| *Anacamptis sancta* (L.) R. M. Bateman | *Glebionis coronaria* (L.) Spach | *Polycarpon tetraphyllum* (L.) L. |
| *Anagallis arvensis* L. | *Hedypnois rhagadioloides* (L.) F. W. Schmidt | *Polygonum equisetiforme* Sm. |
| *Anchusa hybrida* Ten. | *Helichrysum stoechas* (L.) Moench | *Ricinus communis* L.* |
| *Anisantha rigida* (Roth) Hyl. | *Heliotropium hirsutissimum* Grauer | *Rostraria smyrnacea* (Trin.) H. Scholz |
| *Anisantha tectorum* (L.) Nevski | *Hordeum vulgare* L. | *Rumex conglomeratus* Murray |
| *Arundo donax* L. | *Hormuzakia aggregata* (Lehm.) Guşul. | *Sagina apetala* Ard. |
| *Asteriscus aquaticus* (L.) Less. | *Hymenocarpus circinnatus* (L.) Savi | *Sagina maritima* Don |
| *Avena sterilis* L. | *Hyoscyamus albus* L. | *Salvia viridis* L. |
| *Cakile maritima* Scop. | *Lagurus ovatus* L. | *Sarcopoterium spinosum* (L.) Spach |
| *Campanula stellaris* Boiss. | *Lampranthus multiradiatus* (Jacq.) N.E.Br.* | *Senecio × berythaeus* A.Camus & Gomb. |
| *Capparis sicula* Veill. | *Lantana camara* L.* | *Sideritis romana* L. |
| *Cardopatum corymbosum* (L.) Pers. | *Leucaena leucocephala* (Lam.) de Wit* | *Silene aegyptiaca* (L.) L. |
| *Carissa macrocarpa* (Eckl.) A.DC.* | *Limbarda crithmoides* (L.) Dumort. | *Silene colorata* Poir. |
| *Carpobrotus edulis* (L.) N.E.Br.* | *Limonium mousterdei* Domina, Erben & Raimondo | *Silybum Marianum* (L.) Gaertn. |
| *Carthamus tenuis* (Boiss. & C. I. Blanche) Bornm. | *Limonium postii* Domina, Erben & Raimondo | *Sisymbrium officinale* (L.) Scop. |
| *Centaurea procurrens* Spreng. | *Limonium virgatum* (Willd.) Fourr. | *Sonchus oleraceus* L. |
| *Cerastium glomeratum* Thuill. | *Lotus angustissimus* L. | *Sphagneticola trilobata* (L.) Pruski* |
| *Cichorium pumilum* Jacq. | *Lotus cytisoides* L. | *Sporobolus pungens* (Schreb.) Kunth |
| *Convolvulus secundus* Desr. | *Lotus halophilus* Boiss. & Spruner | *Strelitzia reginae* Banks* |
| *Cota palaestina* Kotschy | *Lotus edulis* L. | *Thymbra capitata* (L.) Cav. |
| *Crepis aculeata* (DC.) Boiss. | *Lycopersicon esculentum* Mill.* | *Thymelaea hirsuta* (L.) Endl. |
| *Crepis palaestina* (Boiss.) Bornm. | *Malva oxyloba* Boiss. | *Tordylium trachycarpum* (Boiss.) Al-Eisawi |
| *Crithmum maritimum* L. | *Malva* sp. | *Tragopogon porrifolius* L. |
| *Crucianella aegyptiaca* L. | *Medicago littoralis* Loisel. | *Trifolium glanduliferum* Boiss. |
| *Cuscuta epithymum* (L.) L. | *Mercurialis annua* L. | *Trifolium purpureum* Loisel. |
| *Cyclamen persicum* Mill. | *Ochlopoa annua* (L.) H. Scholz | *Trifolium resupinatum* L. |
| *Cynodon dactylon* (L.) Pers. | *Onobrychis crista-galli* (L.) Lam. | *Trifolium scabrum* L. |
| *Cyperus rotundus* L. | *Orobancha nana* (Reut.) Beck | *Umbilicus intermedius* Boiss. |
| *Dactyloctenium aegyptium* (L.) Willd. | *Oxalis pes-caprae* L.* | *Urospermum picroides* (L.) F. W. Schmidt |
| *Daucus carota* L. | *Pancratium maritimum* L. | *Valantia muralis* L. |
| *Digitaria sanguinalis* (L.) Scop. | *Parapholis incurva* (L.) C. E. Hubb. | *Verbascum sinuatum* L. |
| *Dittrichia viscosa* (L.) Greuter | *Parietaria judaica* L. | *Veronica cymbalaria* Bodard |
| *Echium angustifolium* Mill. | *Paronychia argentea* Lam. | *Washingtonia* sp.* |
| *Elytrigia juncea* (L.) Nevski | *Phagnalon rupestre* (L.) DC. | *Yucca gigantea* Lem.* |
| *Epilobium tetragonum* L. |  |  |

Analysis of floristic data by TWINSPAN clustered the 78 quadrats into 17 quadrat groups labeled F-A to F-Q (Table 3). *M. crassifolia* had the highest constancy and abundance in groups F-D, F-G and F-I. In contrast, groups F-C, F-F, F-K, F-M, F-N, F-O, F-P and F-Q completely excluded this species.

**Table 3.**
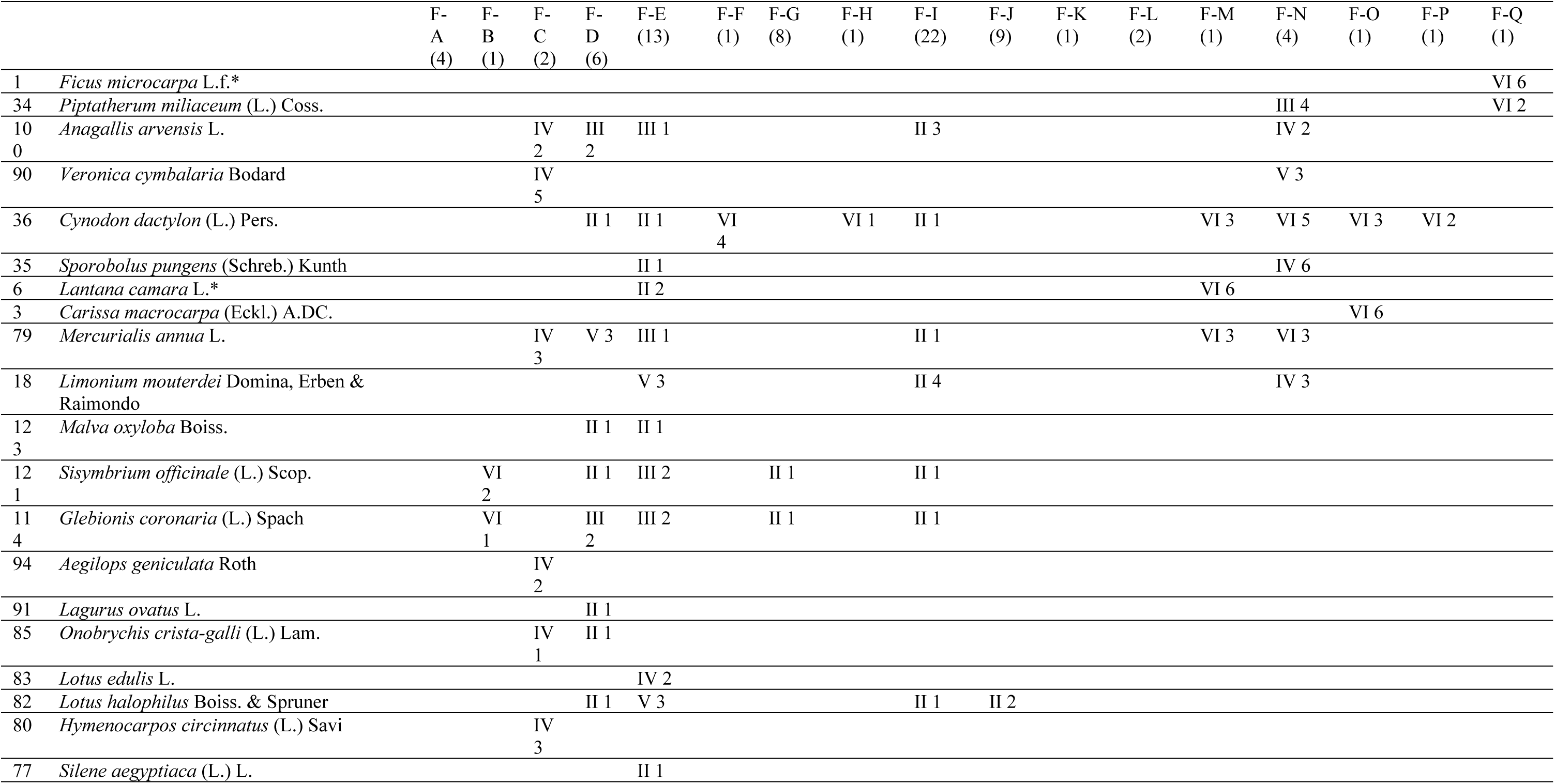
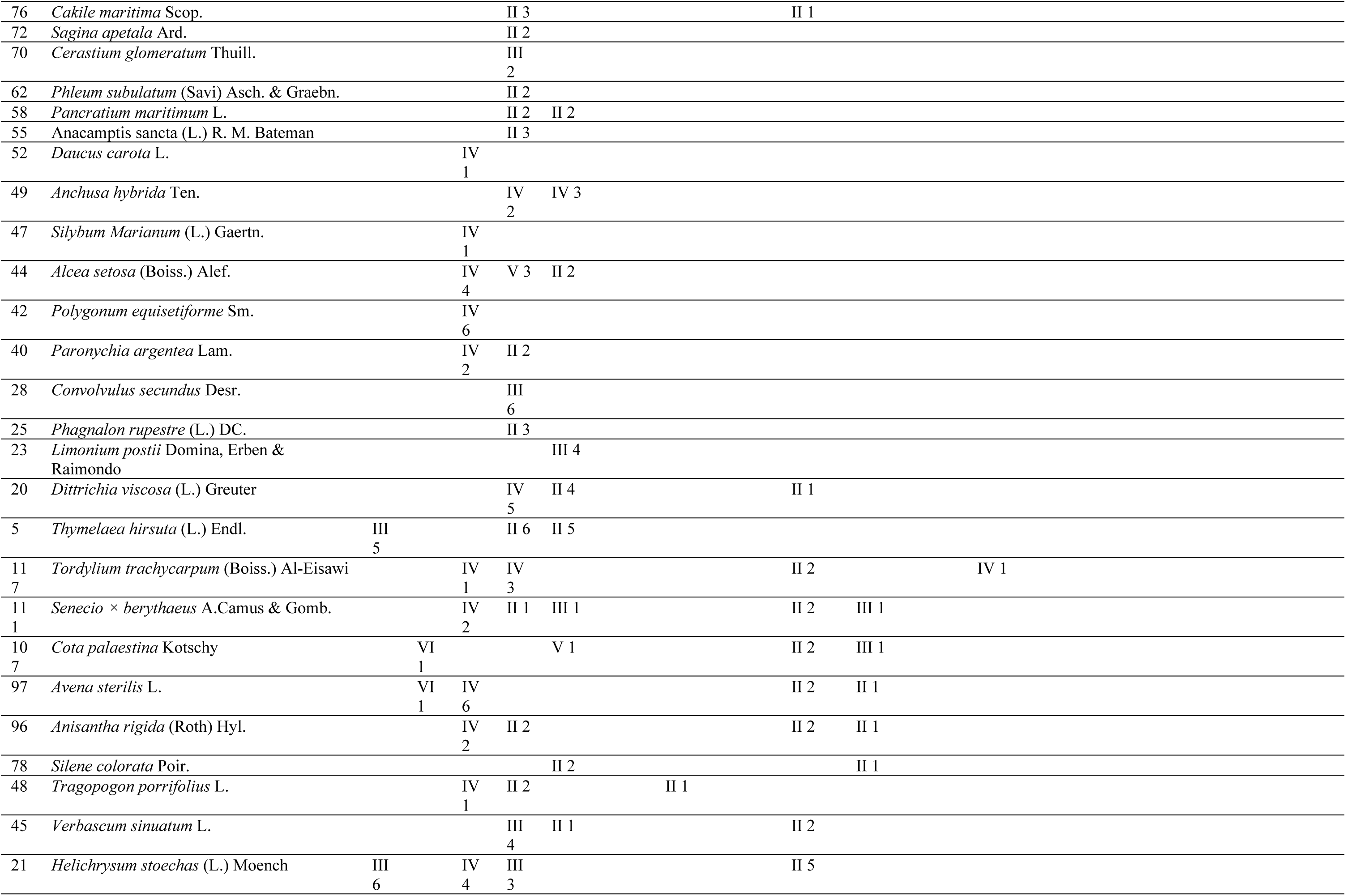
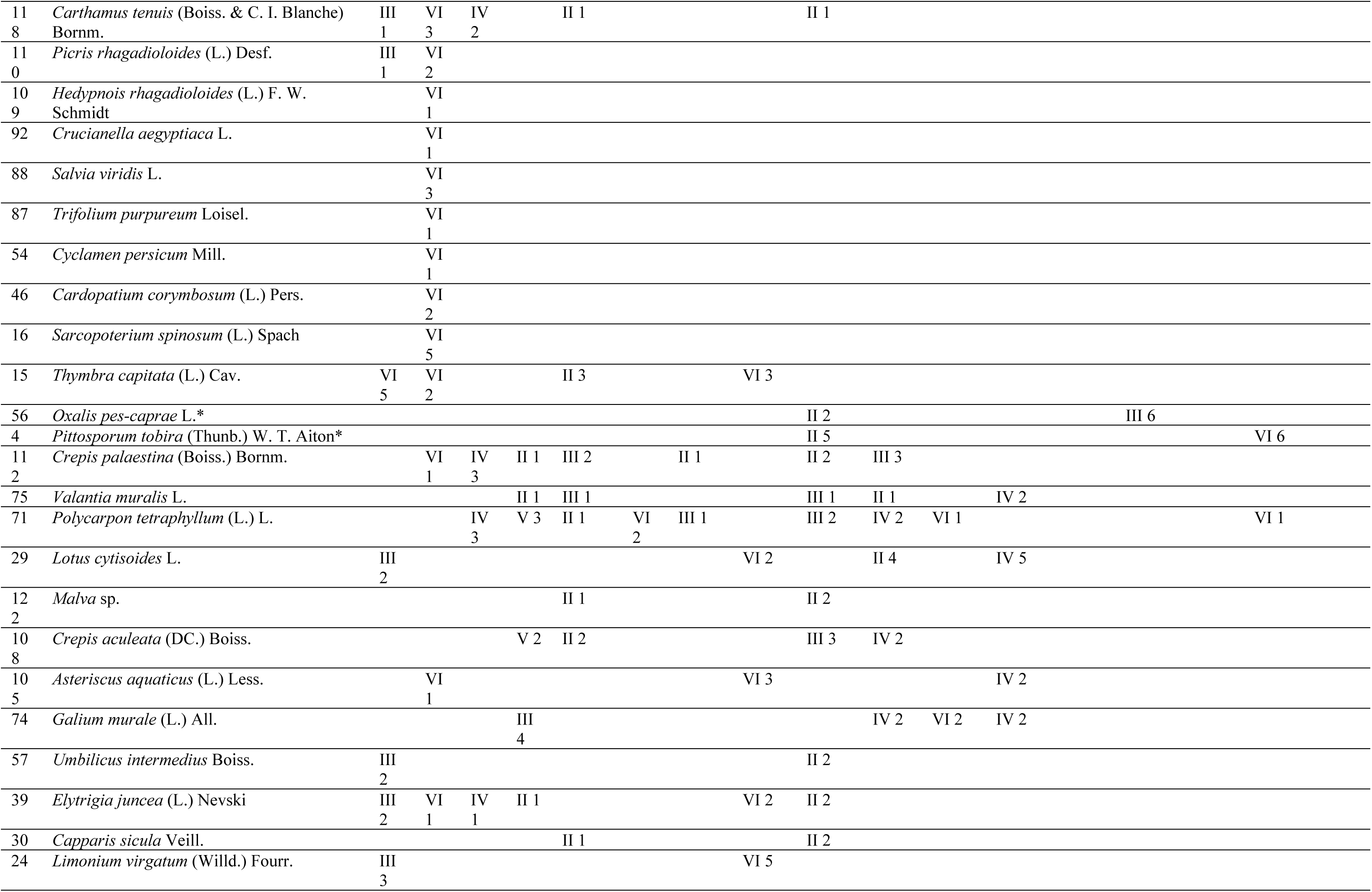
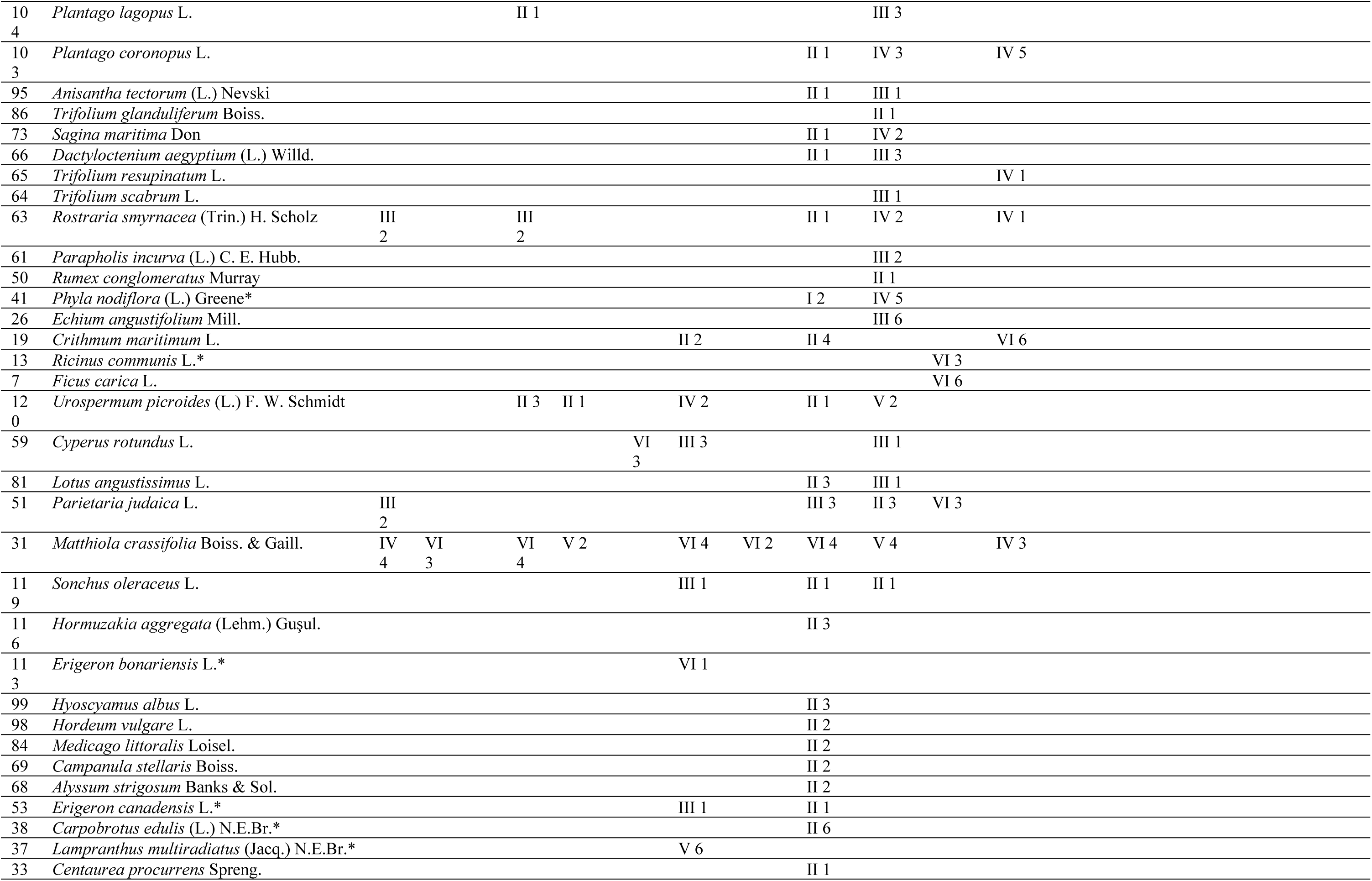
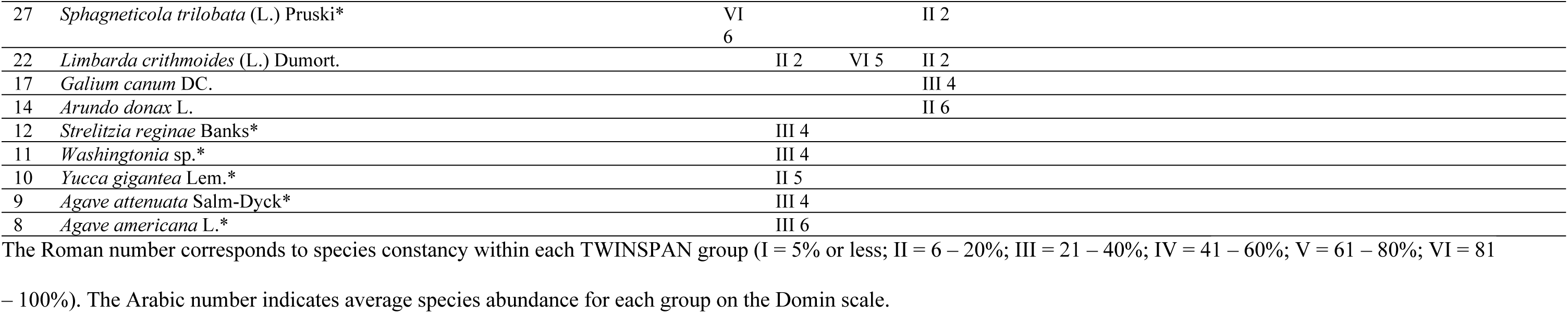
TWINSPAN analysis of floristic data set collected in Ras Beirut (Quadrat groups: F-A to F-Q, (number of quadrats), Alphabetical naming of quadrat groups by floristic and life fom classification are not related.).

The low community similarity, patchy species distribution, and predominance of habitat non-specific species reported by [55] in their study of the floristics of the Lebanese coast was confirmed in this study. High floristic variability between and within different sites resulted in a large number of groups (58.8%) consisting of no more than two quadrats. Only one group (F-E) consisted of a large number of quadrats and represented a perceptible community of sparse vegetation on sandstone outcrops. Other groups were not site specific, but included quadrats exposed to similar disturbance; for example, in group G the nine quadrats were sampled from street medians and side walks and consisted of a combination of evergreen exotic ornamental species such as *Agave americana, A. attenuata and Lampranthus multiradiatus*. Similarly, F-T included quadrats characterized by a high representation of graminoids *Cyperus rotundus* and *Cynodon dactylon* which often grow in gardens and street medians under and around evergreen ornamentals such the shrub *Pittosporum tobira* and the creeping herbaceous forb *Sphagneticola trilobata*.

One problem we encountered with florsitics based TWINSPAN analysis is that many groups did not represent actual communities i.e. plant species found in an area are unique and capable of coexisting as distinct, recognizable units that are repeated regularly in response to biotic and environmental variations [67, 68, 69,70,71]. For example, group F-E, which included about 28% of sampled quadrats, consisted of several distinct vegetation assemblages that occur in different habitats, both semi-natural and anthropogenic, and the target species, a stress-tolerant ruderal, was the only common indicator species between these assemblages.

Life form description of plant species yielded 55 different life forms (Table 4).

**Table 4.**
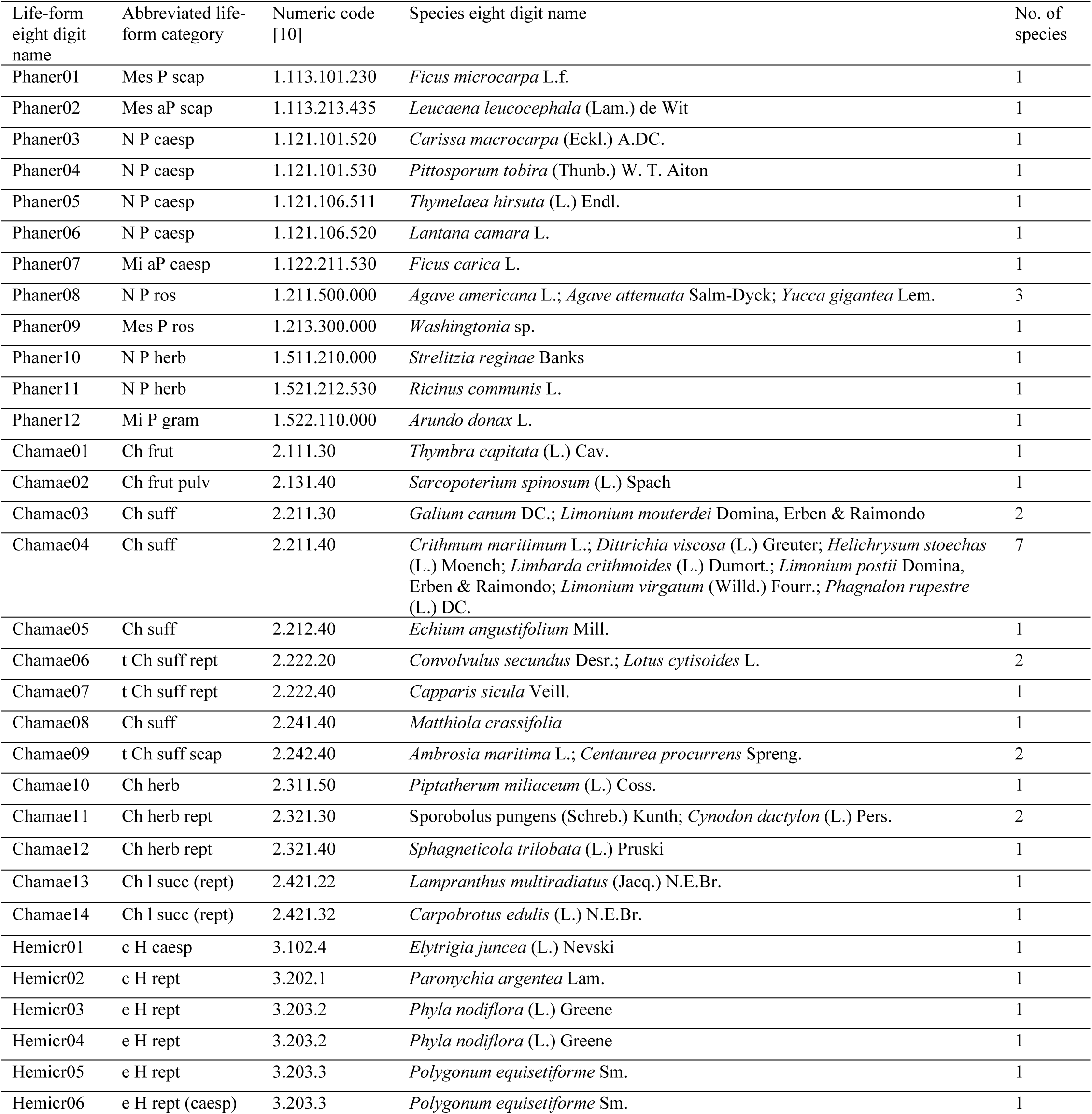
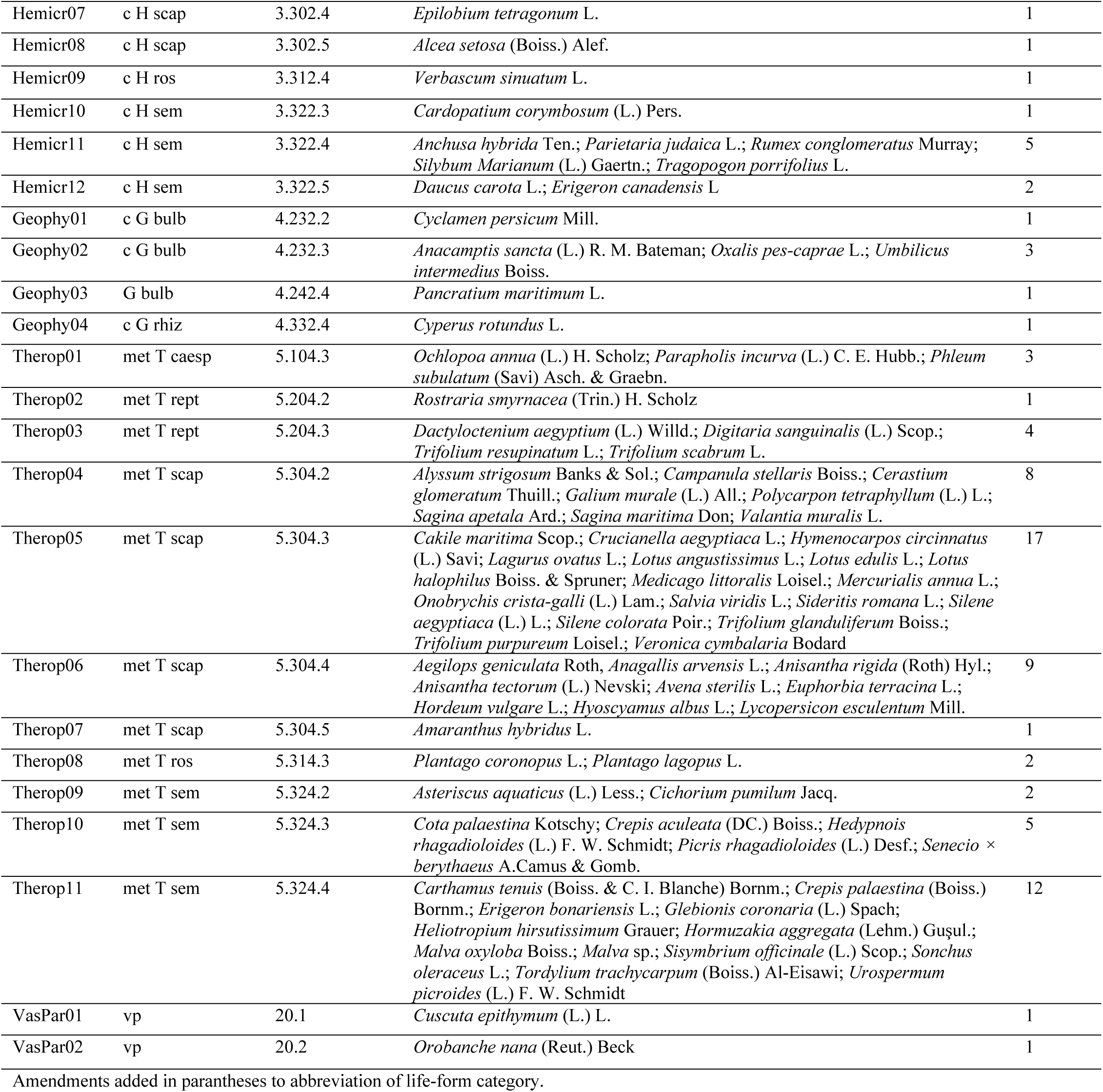
Life form of plant species from 78 quadrats in 12 sites in Ras Beirut.

| Life-form eight digit name | Abbreviated life-form category | Numeric code [10] | Species eight digit name | No. of species |
| --- | --- | --- | --- | --- |
| Phaner01 | Mes P scap | 1.113.101.230 | *Ficus microcarpa* L.f. | 1 |
| Phaner02 | Mes aP scap | 1.113.213.435 | *Leucaena leucocephala* (Lam.) de Wit | 1 |
| Phaner03 | N P caesp | 1.121.101.520 | *Carissa macrocarpa* (Eckl.) A.DC. | 1 |
| Phaner04 | N P caesp | 1.121.101.530 | *Pittosporum tobira* (Thunb.) W. T. Aiton | 1 |
| Phaner05 | N P caesp | 1.121.106.511 | *Thymelaea hirsuta* (L.) Endl. | 1 |
| Phaner06 | N P caesp | 1.121.106.520 | *Lantana camara* L. | 1 |
| Phaner07 | Mi aP caesp | 1.122.211.530 | *Ficus carica* L. | 1 |
| Phaner08 | N P ros | 1.211.500.000 | *Agave americana* L.; *Agave attenuata* Salm-Dyck; *Yucca gigantea* Lem. | 3 |
| Phaner09 | Mes P ros | 1.213.300.000 | *Washingtonia* sp. | 1 |
| Phaner10 | N P herb | 1.511.210.000 | *Strelitzia reginae* Banks | 1 |
| Phaner11 | N P herb | 1.521.212.530 | *Ricinus communis* L. | 1 |
| Phaner12 | Mi P gram | 1.522.110.000 | *Arundo donax* L. | 1 |
| Chamae01 | Ch frut | 2.111.30 | *Thymbra capitata* (L.) Cav. | 1 |
| Chamae02 | Ch frut pulv | 2.131.40 | *Sarcopoterium spinosum* (L.) Spach | 1 |
| Chamae03 | Ch suff | 2.211.30 | *Galium canum* DC.; *Limonium mousterdei* Domina, Erben & Raimondo | 2 |
| Chamae04 | Ch suff | 2.211.40 | *Crithmum maritimum* L.; *Dittrichia viscosa* (L.) Greuter; *Helichrysum stoechas* (L.) Moench; *Limbaria crithmoides* (L.) Dumort.; *Limonium postii* Domina, Erben & Raimondo; *Limonium virgatum* (Willd.) Fourr.; *Phagnalon rupestre* (L.) DC. | 7 |
| Chamae05 | Ch suff | 2.212.40 | *Echium angustifolium* Mill. | 1 |
| Chamae06 | t Ch suff rept | 2.222.20 | *Convolvulus secundus* Desr.; *Lotus cytisoides* L. | 2 |
| Chamae07 | t Ch suff rept | 2.222.40 | *Capparis sicula* Veill. | 1 |
| Chamae08 | Ch suff | 2.241.40 | *Matthiola crassifolia* | 1 |
| Chamae09 | t Ch suff scap | 2.242.40 | *Ambrosia maritima* L.; *Centaurea procurrens* Spreng. | 2 |
| Chamae10 | Ch herb | 2.311.50 | *Piptatherum miliaceum* (L.) Coss. | 1 |
| Chamae11 | Ch herb rept | 2.321.30 | *Sporobolus pungens* (Schreb.) Kunth; *Cynodon dactylon* (L.) Pers. | 2 |
| Chamae12 | Ch herb rept | 2.321.40 | *Sphagneticola trilobata* (L.) Pruski | 1 |
| Chamae13 | Ch l succ (rept) | 2.421.22 | *Lampranthus multiradiatus* (Jacq.) N.E.Br. | 1 |
| Chamae14 | Ch l succ (rept) | 2.421.32 | *Carpobrotus edulis* (L.) N.E.Br. | 1 |
| Hemicr01 | c H caesp | 3.102.4 | *Elytrigia juncea* (L.) Nevski | 1 |
| Hemicr02 | c H rept | 3.202.1 | *Paronychia argentea* Lam. | 1 |
| Hemicr03 | e H rept | 3.203.2 | *Phyla nodiflora* (L.) Greene | 1 |
| Hemicr04 | e H rept | 3.203.2 | *Phyla nodiflora* (L.) Greene | 1 |
| Hemicr05 | e H rept | 3.203.3 | *Polygonum equisetiforme* Sm. | 1 |
| Hemicr06 | e H rept (caesp) | 3.203.3 | *Polygonum equisetiforme* Sm. | 1 |
| Hemicr07 | c H scap | 3.302.4 | *Epilobium tetragonum* L. | 1 |
| Hemicr08 | c H scap | 3.302.5 | *Alcea setosa* (Boiss.) Alef. | 1 |
| Hemicr09 | c H ros | 3.312.4 | *Verbascum sinuatum* L. | 1 |
| Hemicr10 | c H sem | 3.322.3 | *Cardopatum corymbosum* (L.) Pers. | 1 |
| Hemicr11 | c H sem | 3.322.4 | *Anchusa hybrida* Ten.; *Parietaria judaica* L.; *Rumex conglomeratus* Murray;<br>*Silybum Marianum* (L.) Gaertn.; *Tragopogon porrifolius* L. | 5 |
| Hemicr12 | c H sem | 3.322.5 | *Daucus carota* L.; *Erigeron canadensis* L. | 2 |
| Geophy01 | c G bulb | 4.232.2 | *Cyclamen persicum* Mill. | 1 |
| Geophy02 | c G bulb | 4.232.3 | *Anacamptis sancta* (L.) R. M. Bateman; *Oxalis pes-caprae* L.; *Umbilicus intermedius* Boiss. | 3 |
| Geophy03 | G bulb | 4.242.4 | *Pancratium maritimum* L. | 1 |
| Geophy04 | c G rhiz | 4.332.4 | *Cyperus rotundus* L. | 1 |
| Therop01 | met T caesp | 5.104.3 | *Ochlopoa annua* (L.) H. Scholz; *Parapholis incurva* (L.) C. E. Hubb.; *Phleum subulatum* (Savi) Asch. & Graebn. | 3 |
| Therop02 | met T rept | 5.204.2 | *Rostraria smyrnacea* (Trin.) H. Scholz | 1 |
| Therop03 | met T rept | 5.204.3 | *Dactyloctenium aegyptium* (L.) Willd.; *Digitaria sanguinalis* (L.) Scop.;<br>*Trifolium resupinatum* L.; *Trifolium scabrum* L. | 4 |
| Therop04 | met T scap | 5.304.2 | *Alyssum strigosum* Banks & Sol.; *Campanula stellaris* Boiss.; *Cerastium glomeratum* Thuill.; *Galium murale* (L.) All.; *Polycarpon tetraphyllum* (L.) L.;<br>*Sagina apetala* Ard.; *Sagina maritima* Don; *Valantia muralis* L. | 8 |
| Therop05 | met T scap | 5.304.3 | *Cakile maritima* Scop.; *Crucianella aegyptiaca* L.; *Hymenocarpus circinnatus* (L.) Savi; *Lagurus ovatus* L.; *Lotus angustissimus* L.; *Lotus edulis* L.; *Lotus halophilus* Boiss. & Spruner; *Medicago littoralis* Loisel.; *Mercurialis annua* L.;<br>*Onobrychis crista-galli* (L.) Lam.; *Salvia viridis* L.; *Sideritis romana* L.; *Silene aegyptiaca* (L.) L.; *Silene colorata* Poir.; *Trifolium glanduliferum* Boiss.;<br>*Trifolium purpureum* Loisel.; *Veronica cymbalaria* Bodard | 17 |
| Therop06 | met T scap | 5.304.4 | *Aegilops geniculata* Roth; *Anagallis arvensis* L.; *Anisantha rigida* (Roth) Hyl.;<br>*Anisantha tectorum* (L.) Nevski; *Avena sterilis* L.; *Euphorbia terracina* L.;<br>*Hordeum vulgare* L.; *Hyoscyamus albus* L.; *Lycopersicon esculentum* Mill. | 9 |
| Therop07 | met T scap | 5.304.5 | *Amaranthus hybridus* L. | 1 |
| Therop08 | met T ros | 5.314.3 | *Plantago coronopus* L.; *Plantago lagopus* L. | 2 |
| Therop09 | met T sem | 5.324.2 | *Asteriscus aquaticus* (L.) Less.; *Cichorium pumilum* Jacq. | 2 |
| Therop10 | met T sem | 5.324.3 | *Cota palaestina* Kotschy; *Crepis aculeata* (DC.) Boiss.; *Hedypnois rhagadioloides* (L.) F. W. Schmidt; *Picris rhagadioloides* (L.) Desf.; *Senecio × berythaeus* A.Camus & Gomb. | 5 |
| Therop11 | met T sem | 5.324.4 | *Carthamus tenuis* (Boiss. & C. I. Blanche) Bornm.; *Crepis palaestina* (Boiss.) Bornm.; *Erigeron bonariensis* L.; *Glebionis coronaria* (L.) Spach;<br>*Heliotropium hirsutissimum* Grauer; *Hormuzakia aggregata* (Lehm.) Guşul.;<br>*Malva oxyloba* Boiss.; *Malva* sp.; *Sisymbrium officinale* (L.) Scop.; *Sonchus oleraceus* L.; *Tordylium trachycarpum* (Boiss.) Al-Eisawi; *Urospermum picroides* (L.) F. W. Schmidt | 12 |
| VasPar01 | vp | 20.1 | *Cuscuta epithymum* (L.) L. | 1 |
| VasPar02 | vp | 20.2 | *Orobancha nana* (Reut.) Beck | 1 |

Amendments added in parantheses to abbreviation of life-form category.

Results revealed that more than half of all recorded species were therophytes with a total of 64 autotrophic therophyte and two heterotrophic annual vascular parasites. The high representation of therophytes reflects high disturbance of study sites [36]. Fig 1 presents the life-form spectrum defining the associates of *M. crassifolia*. Chamaephytes constituted the most prominent perennial life form and included 24 species. Over half of all chamaephytes were either regional or national endemics and only three were not native. Phanerophytes were represented by 14 species, 10 of which were not native. Perennials characterized by a periodic shoot reduction were represented by 15 hemicryptophytes and six geophytes.

Classification of data according to life form in 11 quadrat groups. *M. crassifolia* was highly represented in two of these groups a percent cover greater than 75% in 81-100% of quadrats within these groups (Table 5). Examples of life forms in these groups include, unbranched dwarf palm like trees (Phaner08), typical and tall evergreen dwarf-shrubs (Chamae03 & Chamae04), low reptant evergreen succulents (Chamae14), tall drought-deciduous hemicryptophytes (Hemicr01) and small reptant evergreen hemicryptophytes (Hemicr03) were common. Ornamental examples of these life forms include *Agave* and *Yucca* species (Phaner08), cultivated Sea Lavender species (Chamae03 and Chamae04), and *Lampranthus multiradiatus* (Chamae13).

**Table 5.** TWINSPAN analysis of Life form data set collected in Ras Beirut (Quadrat groups: L-A to L-J (alphabetical number, (number of quadrats), Alphabetical naming of quadrat groups by floristic and life fom classification are not related.).

|  |  | L-A (1) | L-B (3) | L-C (9) | L-D (12) | L-E (26) | L-F (14) | L-G (4) | L-H (4) | L-I (2) | L-J (2) | L-K (1) |
| --- | --- | --- | --- | --- | --- | --- | --- | --- | --- | --- | --- | --- |
|  |  | - | VI 2 | VI 4 | V 4 | VI 4 | V 2 | IV 3 | - | - | - | - |
| 8 | Phaner08 |  |  | IV 5 |  |  |  |  |  |  |  |  |
| 9 | Phaner09 |  | V 5 | II 1 |  |  |  |  |  |  |  |  |
| 10 | Phaner10 |  | III 5 | II 3 |  |  |  |  |  |  |  |  |
| 25 | Chamae13 |  | III 6 | IV 5 |  |  |  |  |  |  |  |  |
| 38 | Hemicr12 |  | III 1 | II 1 |  | II 1 |  |  |  |  |  |  |
| 31 | Hemicr05 |  |  | II 1 |  | I 2 |  |  |  |  |  |  |
| 17 | Chamae05 |  |  |  | II 6 |  |  |  |  |  |  |  |
| 26 | Chamae14 |  |  |  | II 6 |  |  |  |  |  |  |  |
| 29 | Hemicr03 |  |  |  | III 4 | II 3 |  |  |  |  |  |  |
| 45 | Therop03 |  |  |  | IV 2 | I 1 |  |  |  |  |  |  |
| 50 | Therop08 |  |  | II 1 | V 3 | I 1 |  |  |  |  |  |  |
| 44 | Therop02 |  |  |  | IV 2 | III 1 |  |  |  |  |  |  |
| 7 | Phaner07 |  |  |  |  | I 6 |  |  |  |  |  |  |
| 11 | Phaner11 |  |  |  |  | I 3 |  |  |  |  |  |  |
| 12 | Phaner12 |  |  |  |  | II 6 |  |  |  |  |  |  |
| 14 | Chamae02 |  |  |  |  | I 5 |  |  |  |  |  |  |
| 21 | Chamae09 |  |  |  |  | I 1 |  |  |  |  |  |  |
| 28 | Hemicr02 |  |  |  |  | II 2 |  |  |  |  |  |  |
| 32 | Hemicr06 |  |  |  |  | I 6 |  |  |  |  |  |  |
| 34 | Hemicr08 |  |  |  |  | III 3 | II 1 |  |  |  |  |  |
| 35 | Hemicr09 |  |  |  |  | III 2 |  |  |  |  |  |  |
| 36 | Hemicr10 |  |  |  |  | I 2 |  |  |  |  |  |  |
| 39 | Geophy01 |  |  |  |  | I 2 |  |  |  |  |  |  |
| 48 | Therop06 |  |  |  | III 1 | IV 3 | II 1 |  | III 2 |  |  |  |
| 18 | Chamae06 |  |  |  | III 4 | II 4 |  | III 2 |  |  |  |  |
| 43 | Therop01 | VI 1 |  |  | II 2 | I 2 |  |  |  |  |  |  |
| 46 | Therop04 |  |  | III 1 | VI 3 | IV 2 | III 1 |  |  | VI 2 |  |  |
| 27 | Hemicr01 |  |  | II 1 | II 2 | II 2 |  | III 2 |  |  |  |  |
| 42 | Geophy04 |  |  | III 3 | III 1 |  |  |  |  | IV 3 |  |  |
| 53 | Therop11 |  | V 2 | V 2 | V 2 | IV 3 | IV 1 |  |  |  |  |  |
| 20 | **Chamae08** |  | **VI 2** | **VI 4** | **V 4** | **VI 4** | **V 2** | **IV 3** |  |  |  |  |
| 37 | Hemicr11 |  |  |  | II 1 | IV 2 | III 2 |  |  |  |  |  |
| 52 | Therop10 |  |  |  | VI 2 | IV 2 | IV 1 |  |  |  |  |  |
| 4 | Phaner04 |  |  |  | II 6 | I 3 |  |  |  | IV 6 |  |  |
| 5 | Phaner05 |  |  |  |  | II 5 | II 5 |  |  |  |  |  |
| 13 | Chamae01 |  |  |  | III 4 | II 4 | II 3 | IV 4 |  |  |  |  |
| 16 | Chamae04 |  | V 4 |  | II 4 | III 3 | III 3 | VI 6 |  |  |  |  |
| 19 | Chamae07 |  |  |  |  | I 2 | II 1 |  |  |  |  |  |
| 40 | Geophy02 |  |  |  |  | III 2 | II 1 |  | III 6 |  |  |  |
| 47 | Therop05 |  |  |  | IV 1 | IV 3 | V 3 | III 3 | VI 4 |  |  |  |
| 51 | Therop09 |  |  |  | II 2 | I 1 |  | III 3 |  |  |  |  |
| 15 | Chamae03 |  |  |  |  | I 3 | VI 4 | III 2 | III 3 |  |  |  |
| 24 | Chamae12 |  |  |  |  | I 2 | II 1 |  |  | IV 6 |  |  |
| 41 | Geophy03 |  |  |  |  |  | II 2 |  |  |  |  |  |
| 3 | Phaner03 |  |  |  |  |  |  |  |  |  | IV 6 |  |
| 6 | Phaner06 |  |  |  |  |  | II 2 |  |  |  | IV 6 |  |
| 23 | Chamae11 |  |  |  |  | I 1 | II 1 | III 2 | VI 6 | VI 3 | VI 3 |  |
| 30 | Hemicr04 | VI 6 |  |  |  |  |  |  |  |  |  |  |

Five groups excluded the target species and the dominant life form in these groups was mostly phanerophytes. These include mesophyllous large evergreen trees with spherical crown restricted to their upper half (Phaner01), mesophyllous normal-sized evergreen shrubs with spherical crown extending to near their base (Phaner04), microphyllous normal-sized evergreen shrubs with spherical crown extending to near their base (Phaner03), and mesophyllous tall deciduous shrub with spherical crown extending to near the base of the shrub (Phaner07). Ornamental examples of these life forms include various shade trees (Phaner01), and shrubs used as hedges such as *Pittosporum tobira* (Phaner04 and Phaner03).

Other groups that excluded the target species consisted mostly of typical evergreen reptant herbaceous chamaephytes (Chamae12). Ornamental plant species belonging to this life form and similar life forms include turfgrass species and the Singapore Daisy, *Sphagneticola trilobata*.

In Table 6 below, we integrated floristic and life-form classification results along into a single matrix by identifying common quadrats intersecting both classifications. Using this stepwise approach we generated a new set of quadrat groups which included quadrats that shared similar life form and species composition. To assess the relevance of these newly generated groups to *M. crassifolia* prevalence, we calculated constancy and abundance of *M. crassifolia* within each group. This stepwise approach generated 30 quadrat groups, 8 which were highly favorable to *M. crassifolia*, and 12 which excluded it. We then proceeded to describe life form and species prevalent in these groups.

**Table 6.**
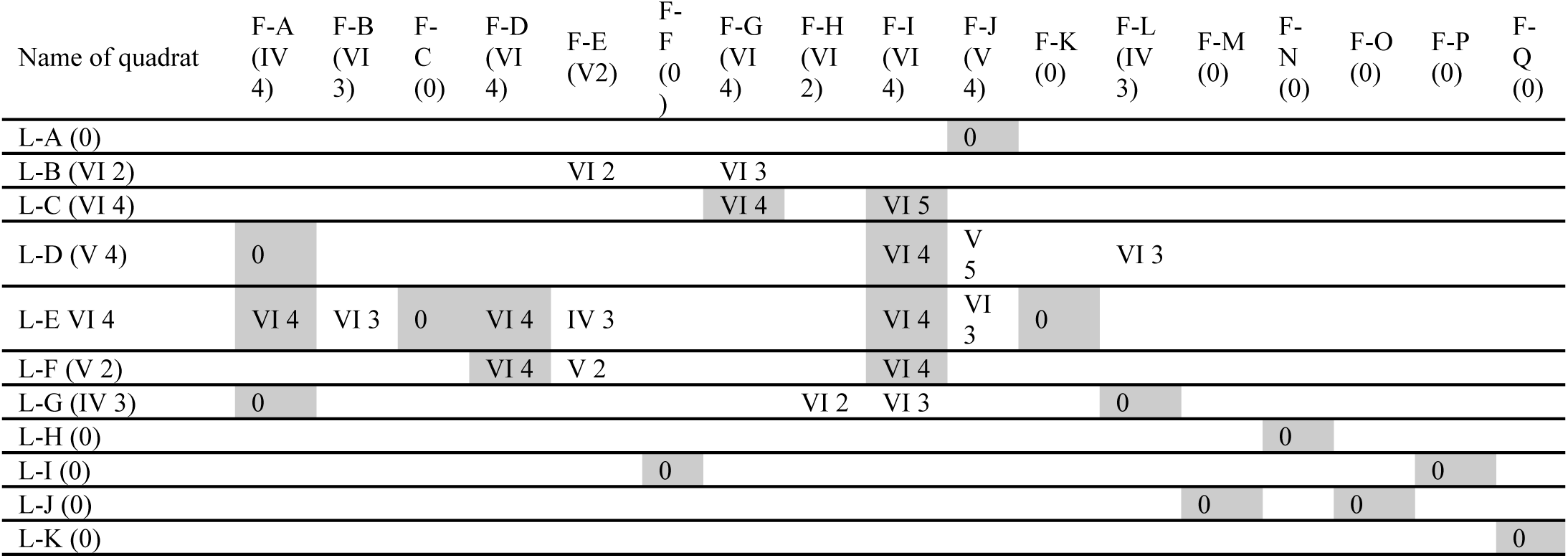
Matrix of floristic and life-form classifications of quadrats from plant data set collected in Ras Beirut. Intersections show favorable and unfavorable vegetation assemblages for *M. crassifolia* represented by constancy and abundance. (Quadrat groups: L-A to L-J and F-A to F-Q, F = floristic, L=life form (Alphabetical naming of quadrat groups by floristic and life fom classification are not related), constancy (I = 5% or less; II = 6 – 20%; III = 21 – 40%; IV = 41 – 60%; V = 61 – 80%; VI = 81 – 100%), average cover (1-5)).

Quadrat groups of high representation of target species formed by the intersection of both floristic and life form data classifications are listed in Table 7. The intersections that resulted in quadrat groups with the highest representation of the target species belonged to 4 out of 11 quadrat groups that were derived from the classification of the life form data set (L-C, L-D, L-E and L-F) and 4 out of 17 quadrat groups that were derived from the classification of the floristic data set (F-A, F-D, F-G and F-I).

**Table 7.** Urban plant habitat analogs in Beirut for M. crassifolia resulting from quadrat groups of high representation of target species following a stepwise approach that intersects floristic and life form data classifications.

| Floristic Classification | Life form Classification | Average Cover of Target Species | Constancy of Target Species | Description of urban habitat analog: life form | Description of urban habitat analog: Plant habitat and species |
| --- | --- | --- | --- | --- | --- |
| I | C | 5 | VI | Quadrat groups dominated solely by suffruticose chamaephytes, the life form of the target species, sometimes including fruticose chamaephytes and caespitose nanophanerophytes with scale like leaves | The highest representation of the target species was only revealed through the matrix. The quadrat group shows that the target species probably prefers to be alone. |
| A | E | 4 | VI |  | Species poor quadrat group. *Matthiola crassifolia* was the only species consistently common between the quadrats. Perennials that less significantly occurred included *Thymra capitata* and *Thymelaea hirsuta*. |
| G | C | 4 | VI | Mostly quadrat groups describing vegetation of street medians. Low lying spreading succulent chamaephytes growing spontaneously or used as ground cover, sometimes interspersed by rosulate nanophanerophytes. Semi-rosette therophytes behaved as consistent ruderals. | Dominated by palm-like species of Agave and Yucca. *Lampranthus multiradiatus* used as ground cover. Several annuals, most notably *Urospermum picroides*, and *Matthiola crassifolia* behaved as ruderals. |
| I | D | 4 | VI |  | *Polycarpon tetraphyllum* and *Crepis aculeata* were common ruderals - besides *Matthiola crassifolia*. *Carpobrotus edulis* dominated - *Pittosporum tobira* dominated once, but in that case, its canopy was disturbed. |
| D | E | 4 | VI | Very tall Drought-deciduous scapose hemicryptophytes, small and very tall scapose therophytes were consistent ephemeral elements of this quadrat group. | Sandy soil with small rock fragments sometimes alternatively dominated by *Dittrichia viscosa*, *Thymaleae hirsuta* or *Convulvulus secundus*, among other perennials and annuals, but consistently including the target species as well as *Alcea setosa* |
| I | E | 4 | VI | Drought deciduous semi-rosette scapose hemicryptophytes and tall scapose therophytes were regular features in this group of quadrats. Besides graminoid phanerophytes being seldom present as evergreen perennial elements, tall scapose suffrutescent chamaephytes were consistently present at relatively high abundance. | This quadrat group included both anthropogenic and disturbed semi-natural habitats. Sparse vegetation composed of evergreen ornamentals and ruderals growing on a mostly bare sandy soil mixed with gravel in a managed street median or cracks in concrete. Vegetation growing on slightly stabilized sands of a sandy beach; Meeting line of sandstone formation with pedestrian path, abandoned dump site; Mostly bare ground on wet sandstone cliff occupied by sparse vegetation; Mostly bare ground on wet sandstone cliff occupied by sparse vegetation; Part of steep sandstone cliff dominated by *Galium canum*; Sandy soil with small rock fragments and cement dominated with *Arundo donax* |
| I | F | 4 | VI | Typical or tall caespitose and tall scapose suffrutescent chamaephytes codominating vegetation. | Crack in concrete through which few perennial species grow; A bolder protruding from a sandstone cliff allowing for both *Limonium mouterdei* and *Matthiola crassifolia* to grow on it; Part of steep sandstone cliff dominated by *Galium canum* |
| D | F | 4 | VI |  | *Dittrichia viscosa* and *Matthiola crassifolia* dominating vegetation growing on slightly stabilized sands of a sandy beach |

Quadrat groups that excluded *Matthiola crassifolia* formed by the intersection of both floristic and life form data classifications are listed in Table 8. The intersections that resulted in quadrat groups with the highest representation of the target species belonged to 8 out of 11 quadrat groups that were derived from the classification of the life form data set and 11 out of 17 quadrat groups that were derived from the classification of the floristic data set.

**Table 8.**
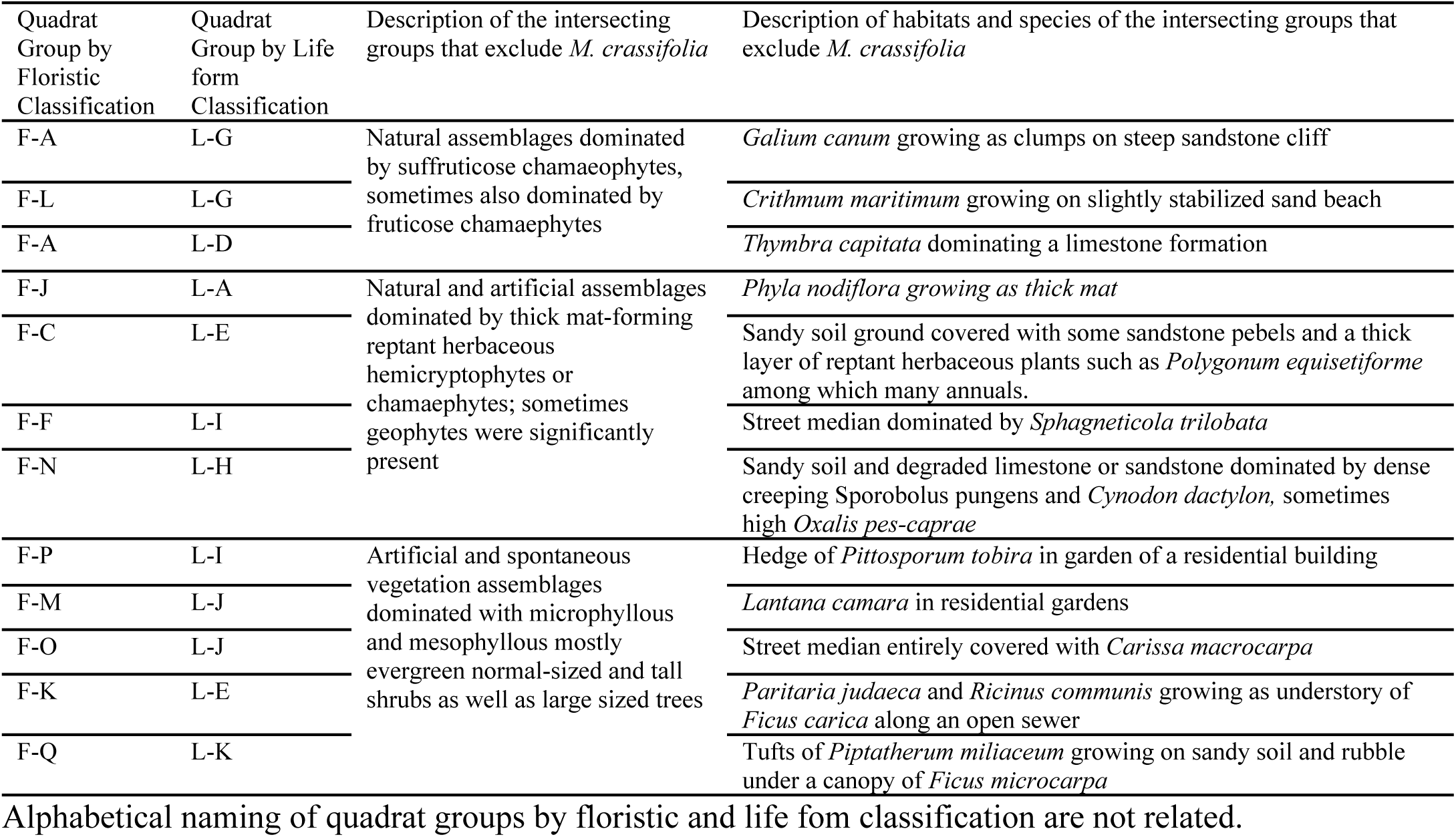
Quadrat groups that excluded target species formed by the intersection of both floristic and life form data classifications.

## Discussion

Floristic surveys are one of two main vegetation description methods used to obtain baseline data on native species of conservation interest and to generate community classification schemes and structure patterns that vary predictably in response to external factors such as environmental stress and disturbance (Table 1). Floristic method uses taxonomic identification and species abundance to describe vegetation. From the perspective of floristics, plant species found in an area are unique and capable of coexisting as distinct, recognizable units that are repeated regularly in response to biotic and environmental variations [67, 68, 69, 70, 71]. The other method, physiognomy, describes vegetation according to external morphology, life form, stratification, and size of each species. There is a consensus that physiognomic and physiological characteristics of plants, including species life-history strategies and population biology, are also important descriptors of vegetation communities [72, 36, 73, 74, 75, 76, 77, 78, 79]. Using either of these methods is a basic step necessary for understanding optimal plant habitats for species of conservation interest [18]. For example, plant communities were characterized to determine suitable habitats for rare species, e.g., [58] and for ecologically and economically important species [80]. Such studies, however, are mostly conducted in natural habitats, and in many instances, deliberately exclude disturbed areas from sampling [80]. In cities, plant habitats are disturbed, and vegetation communities often remain at early successional stages due to this disturbance. Mediterranean cities, where plant diversity and endemism are high, can offer a prospective refuge regardless of whether urban plant habitats are semi-natural or anthropogenic [81]. Plant diversity in Mediterranean cities has been assessed through floristic surveys in Greece, Italy and Spain [82, 83, 84, 85, 86, 87, 88]. The impact of Mediterranean cities on this diversity can be estimated as an extinction debt explained by the city’s current proportion of urban native vegetation and its historical development [57].

One aspect of urban vegetation that challenges field data analysis is the abundance of ruderal plant species which benefit from the absence of interspecific competition that would otherwise occur in later successional stages and colonize bare and disturbed land [36]. By spreading from nearby semi-natural vegetation, ruderals contribute to high variability in urban plant diversity, even between close sites, limiting the value of vegetation description using floristic methods [18]. Some of these ruderal species may be distantly related to agricultural weeds and others to plant species found across transportation networks [24]. The similarity in the infrastructure of a city may explain homogeneity of these urban ruderal species, which out-compete sown species [89]. For example, a 30-year green roof study concluded that spontaneous colonization should be accepted and considered as a design factor; and regional plant communities could serve as a model for seed recruitment and installations [89]. Ruderals are also populating green walls in cities [90]. The peculiarity of our study is that, not only is data analysis influenced by ruderals but the species of conservation interest *M. crassifolia* also behaves as a ruderal. Considering the diversity of habitats the species of conservation interest occupies, it was not possible to resolve this lack of location specificity with floristic assessments, which in turn did not allow us to develop an understanding of urban habitat analogs. Instead, the number of quadrat groups generated by the floristic analysis was large, and some of these clusters did not represent actual plant community assemblages.

Classifying life form data by including percent cover for each category helped specify which life forms and their respective abundance were positively or negatively associated with *M. crassifolia*. Our findings are in line with Kent [18], who emphasized that physiognomy might be more useful as a tool than floristics in highly modified habitats at different scales due to the responses of plant species to macro- and micro-climate conditions. Life history and life form are stronger predictors of underlying population processes than native status and can help explain allelopathic potentials [91, 92].

By using a stepwise approach which combines the two methods, floristics and physiognomy, we were able to minimize the masking effect of ruderal species and to identify life form similarities within distinct vegetation assemblages. In the last decade, researchers have combined life form and floristic vegetation description methods to overcome difficulties in analyzing data in disturbed habitats. For example, Vestergaard [93] generated quadrat groups based on floristic data through TWINSPAN and then described the life-form spectra in each to investigate the relationship between plant diversity and artificial dune development processes. Although similar to our methodology, Vestergaard did not use this combined methodology to define habitat analogs for target plant species. In 2014, a new vegetation classification approach that relies on both physiognomy and floristics over large areas was published under the name EcoVeg [19]. Our approach, however, differs from EcoVeg in that we first mathematically classify physiognomic data and later sort the classifications according to a specific floristic trend. In addition, we base our study on field data collected from small urban habitat sites while EcoVeg uses map data and is meant to classify vegetation over large natural landscapes. On the other hand, our stepwise approach can be integrated as a potential field verification tool with a recent technique proposed by [94] “light detection and ranging (LiDAR) data and model selection techniques.” LiDAR was developed to facilitate the management of urban vegetation for biodiversity conservation by determining potential locations for habitat analogs in cities through the relationship between the extent and vertical structure of urban vegetation.

The information we generated using a stepwise approach integrating floristics and physiognomy, may serve as blueprints for planting designs; it offers a plant selection palette that is not restrictive and does not enforce a native only policy. The urban habitat analogs that we identified include green spaces dominated by palms, low-lying succulents, or shrubs with scale-like leaves. In contrast, the species does not seem to persist in green spaces dominated by turf grass, canopy trees, or vegetation that produces a significant litter.

Furthermore, since knowledge of a target species’ preferred physiognomies includes an understanding of its position in the vertical stratification of its ecological community [18], we were able to identify additional habitats suitable for the introduction of *M. crassifolia*. Our findings revealed that the species could also thrive as part of the low shrub layer under taller nanophyllous shrubs like the Shaggy sparrow-wort, *Thymalea hirsuta*, in the understory of tuft-trees like the fan palm, *Washingtonia robusta*, and within groves of the giant reed, *Arundo donax*. Species belonging to these life forms, or similar ones, dominate many sites in Beirut including street medians and could serve as favorable habitats for *M. crassifolia*. Our findings also show that some exotic invasive species impacted *M. crassifolia* positively. *M. crassifolia* grew in sites dominated by *Carpobrotus edulis*, a potentially invasive in Lebanon, planted at the edge of pedestrian paths. Pedestrians avoided stepping onto these areas, maybe due to their appreciation of *C. edulis* as an evergreen ground cover [95]. As a result, this plant assemblage protected *M. crassifolia* and allowed *C. edulis* to spread constrained by water availability. Removal of invasive plant species should be determined based on its impact on endemic and rare vegetation present in a given region, and eradication should focus on those invasive species that compete with endemic species in general and those of conservation interest especially [96]. Huenneke and Thomson [97] suggest criteria for determining whether such species pose problems for specific rare native taxa and indicated the possibility that some species may be beneficial to endemics. Equipped with the findings above, landscape designers, architects, and managers can better reconcile between desired conservation targets and, socio-behavioral, and aesthetic outcomes by including *M. crassifolia* in an aesthetically pleasing setting. They can design urban habitat analogs that promote the persistence of *M. crassifolia* by excluding from the plant palette native or non-native species belonging to life forms associated with its low representation as reported in this study. Alternatively, they can design an urban habitat analog using a vegetation architecture conducive to the persistence of *M. crassifolia*. In the case established green spaces, they can manage the space to become suitable for *M. crassifolia* by selectively removing species with a life form that is incompatible or that restricts its abundance. In some situations, horticultural techniques, such as pruning, can modify the micro environment without changing species existing on site, to create suitable urban habitat analogs; for example, improving light conditions in cases where species of conservation interest is shade intolerant.

## Conclusion

Given the rate of expansion of urban landscapes [98, 99, 100, 101], increasing species’ site area in a city is highly desired [4]. Our findings can serve as guidance on how to create or modify, through landscape planting designs, suitable habitats for species of conservation interest. By understanding the physiognomy and structure, and environmental conditions in which a species occurs, green areas may be designed to suit the requirements of a target species while established areas may be surveyed for candidate sites suited for the introduction of a target species. Our stepwise approach offers a detailed field assessment tool for urban plant habitat analog characterization.

## Acknowledgements

This paper is derived from the dissertation submitted by M. Itani in partial fulfillment of the requirements for the MSc. degree at the American University of Beirut. We thank Drs. R. Zurayk, N. Farajalla, and K. Knio for their inputs throughout the study. We thank K. Mohamed, A. Jammool, S. El Masri, O. El Tal, R. Atallah and N. Halabi for their in field data collection.

---

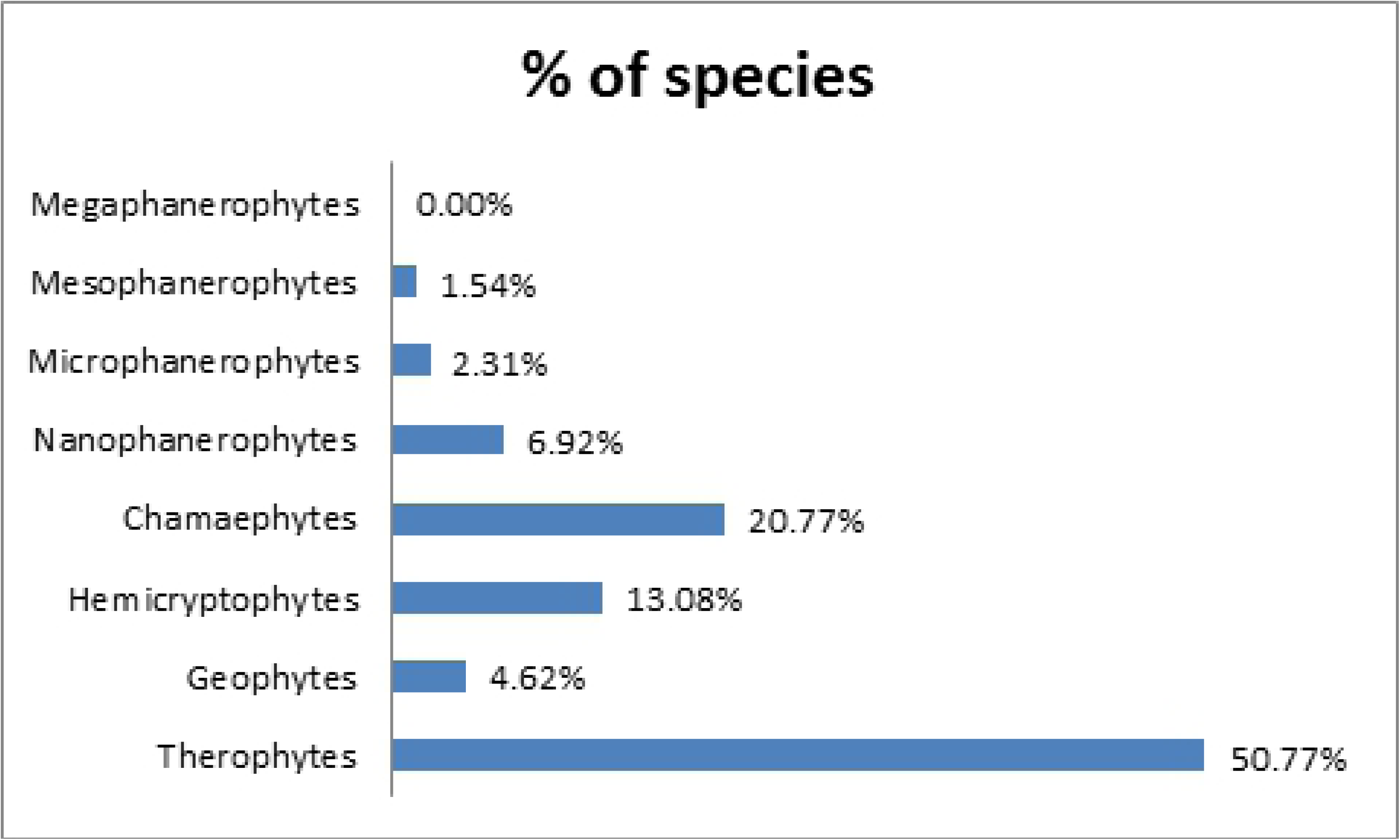

